# The immune vulnerability landscape of the 2019 Novel Coronavirus, SARS-CoV-2

**DOI:** 10.1101/2020.02.08.939553

**Authors:** James Zhu, Jiwoong Kim, Xue Xiao, Yunguan Wang, Danni Luo, Shuang Jiang, Ran Chen, Lin Xu, He Zhang, Lenny Moise, Andres H. Gutierrez, Anne S. De Groot, Guanghua Xiao, John W. Schoggins, Xiaowei Zhan, Tao Wang, Yang Xie

**Author notes:** Co-first authors (James Zhu, Jiwoong Kim, Xue Xiao, Yunguan Wang). Corresponding authors: (1) Tao Wang, Ph.D., Quantitative Biomedical Research Center, Department of Population and Data Sciences, UT Southwestern Medical Center, Dallas, TX, 75390, USA; (2) Yang Xie, Ph.D., Quantitative Biomedical Research Center, Department of Population and Data Sciences, UT Southwestern Medical Center, Dallas, TX, 75390, USA.

## Abstract

The outbreak of the 2019 Novel Coronavirus (SARS-CoV-2) rapidly spread from Wuhan, China to more than 150 countries, areas, or territories, causing staggering numbers of infections and deaths. In this study, bioinformatics analyses were performed on 5,568 complete genomes of SARS-CoV-2 virus to predict the T cell and B cell immunogenic epitopes of all viral proteins, which formed a systematic immune vulnerability landscape of SARS-CoV-2. The immune vulnerability and genetic variation profiles of SARS-CoV were compared with those of SARS-CoV and MERS-CoV. In addition, a web portal was developed to broadly share the data and results as a resource for the research community. Using this resource, we showed that genetic variations in SARS-CoV-2 are associated with loss of B cell immunogenicity, an increase in CD4^+^ T cell immunogenicity, and a minimum loss in CD8^+^ T cell immunogenicity, indicating the existence of a curious correlation between SARS-CoV-2 genetic evolutions and the immunity pressure from the host. Overall, we present an immunological resource for SARS-CoV-2 that could promote both therapeutic/vaccine development and mechanistic research.

## INTRODUCTION

In December 2019, an outbreak of a novel coronavirus (SARS-CoV-2) was reported in Wuhan, China (Li et al., 2020). SARS-CoV-2 rapidly spread to more than 150 other countries, at a speed that is much higher than the Severe Acute Respiratory Syndrome coronavirus (SARS-CoV) and the Middle East Respiratory Syndrome coronavirus (MERS-CoV) (de Wit et al., 2016). Despite the lower mortality rate of SARS-CoV-2 compared with those of SARS-CoV and MERS-CoV, the scale of the SARS-CoV-2 contagion has already caused many more casualties than either of the previous outbreaks. Early research into SARS-CoV-2 has mostly described its epidemiological features (Huang et al., 2020; Li et al., 2020), case reports (Holshue et al., 2020), structural characterizations (Wrapp et al., 2020), basic genomics features (Lu et al., 2020a; Wu et al., 2020; Zhou et al., 2020), *etc*.

Comprehensive profiling of the immunological features of SARS-CoV-2 could broaden our understanding of how this virus interacts with its host. Such analyses could also inform antiviral of immuno-therapeutic development. Antibodies can neutralize viral infectivity in a number of ways, such as interfering with binding to receptors, blocking uptake into cells, *etc*. For SARS-CoV, the human ACE-2 protein is the functional receptor, and the anti-ACE2 antibody can block viral replication (Li et al., 2003). In addition, previous studies have indicated a crucial role of both CD8^+^ and CD4^+^ T cells in SARS-CoV clearance (Chen et al., 2010; Janice Oh et al., 2012), while Janice Oh *et al* also observed that development of SARS-CoV specific neutralizing antibodies requires CD4^+^ T helper cells (Janice Oh et al., 2012).

The investigation of the genetic variations in SARS-CoV-2 over time and the association between the genetic variation and the changes in the immunogenic viral epitopes will provide novel biological insights into the evolution of the virus. In this study, we performed a bioinformatics analysis on 5,568 complete genomes of SARS-CoV-2 virus (from the beginning of the outbreak to May 13^th^, 2020) to profile the class I and class II MHC binding potentials of the SARS-CoV-2 proteins, and also the potential of linear epitopes of the viral proteins to induce antibodies. These data formed the immune vulnerability map of SARS-CoV-2, which was correlated with the genomic variants found in the SARS-CoV-2 strains. The comparison was made against SARS-CoV and MERS-CoV, leading to several interesting observations. All analyses results were made publicly available as a resource to the research community, in the form of the SARS-CoV-2 Immune Viewer: https://qbrc.swmed.edu/projects/2019ncov_immuneviewer/.

## RESULTS

### T and B cell-mediated immune vulnerability landscape of SARS-CoV-2

To explore the immune vulnerability landscape of SARS-CoV-2, we used the NetMHCpan (v4.0) (Jurtz et al., 2017) and NetMHCIIpan (v3.2) (Jensen et al., 2018) software to predict the MHC class I and class II binding peptides of all SARS-CoV-2 proteins (**Fig. 1a**). As there are currently no well-accepted tools for predicting the potential of both class I and II epitopes to induce T cell responses, we used the binding affinity of epitopes to MHCs as a surrogate for epitope immunogenicity, since such binding affinity has been shown to be one of the most important predictors of epitope immunogenicity (Lu et al., 2020b). As this virus was initially reported in China, the counts of the MHC binders for the reference genome were weighted by the allele frequencies of the Chinese population. The results show that there are a small number of genomic regions that showed high peaks of immunogenicity corresponding to a large number of MHC binders in a small neighborhood (60 nt), which could be better potential vaccine targets (**Sup. Table 1**). Of the top 50 MHC class I and MHC class II epitopes, 18 and 6 have already been validated experimentally and documented on the Immune Epitope Database (https://www.iedb.org/), respectively (**Sup. Table 1**). The MHC binding peptide profiles of several other racial populations: US European Caucasian, African American, Hispanic (South or Central American), and Middle Eastern/North African, were also investigated. There are minor differences in the overall immunogenicity of SARS-CoV-2 in different racial populations (**Sup. Fig. 1** and **Sup. Fig. 2**); however, much larger differences can be seen when examining individual HLA alleles (**Fig. 1b**), suggesting a large degree of variance in susceptibility to this virus could exist among different individuals (with different sets of HLA alleles). This suggests that T cell epitope-based therapeutic interventions should focus on genomic regions of the virus that are broadly presentable by as many HLA alleles as possible. This might also underlie the vast degree of differences in clinical manifestations, from being asymptotic to severe complications, when patients are infected with SARS-CoV-2.

**Fig. 1.**
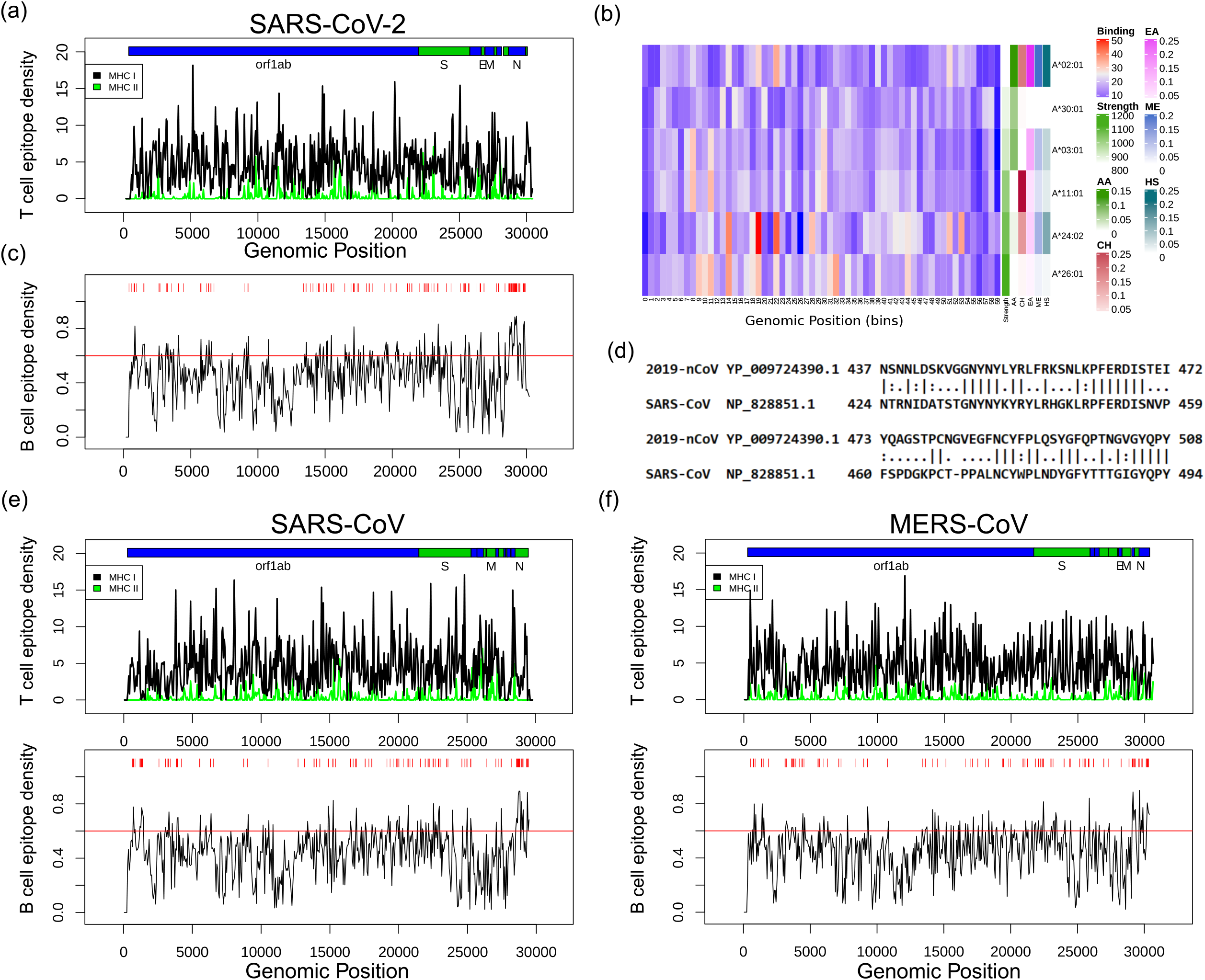
T cell- and B cell-mediated immune vulnerability landscape of SARS-CoV-2. (a) The CD4^+^ and CD8^+^ T cell epitope profiles of SARS-CoV-2. The Y-axis shows the immunogenicity intensity as described in the method section. The X-axis is the genomic coordinate, but with a small number of gaps added by multi-alignment. This is the same for all genomic coordinates shown in this paper. The HLA allele frequency used is that of the Chinese population. (b) Allele-specific T cell epitope profiles for SARS-CoV-2, comparing the different populations. The heatmap represents the number of immunogenic binding epitopes across the binned genomes (500bp) of SARS-CoV-2 for several major HLA-A alleles shown as examples in the racial populations investigated. On the right, the band of “strength” represents the cumulative number of immunogenic peptides, and the other colored bands indicate the HLA allele frequencies for each population. (c) The B cell epitope profiles of the SARS-CoV-2. The Y-axis shows the predicted B cell epitope score. X-axis shows the genomic coordinate. Only showing residues with predicted epitope score >0.6. (d) BLASTing the motif binding domain of the SARS-CoV-2 S protein and the SARS-CoV S protein. (e) The T cell and B cell epitope profiles of SARS-CoV. (f) The T cell and B cell epitope profiles of MERS-CoV. (e) and (f) were both for the Chinese population.

**Fig. 2.**
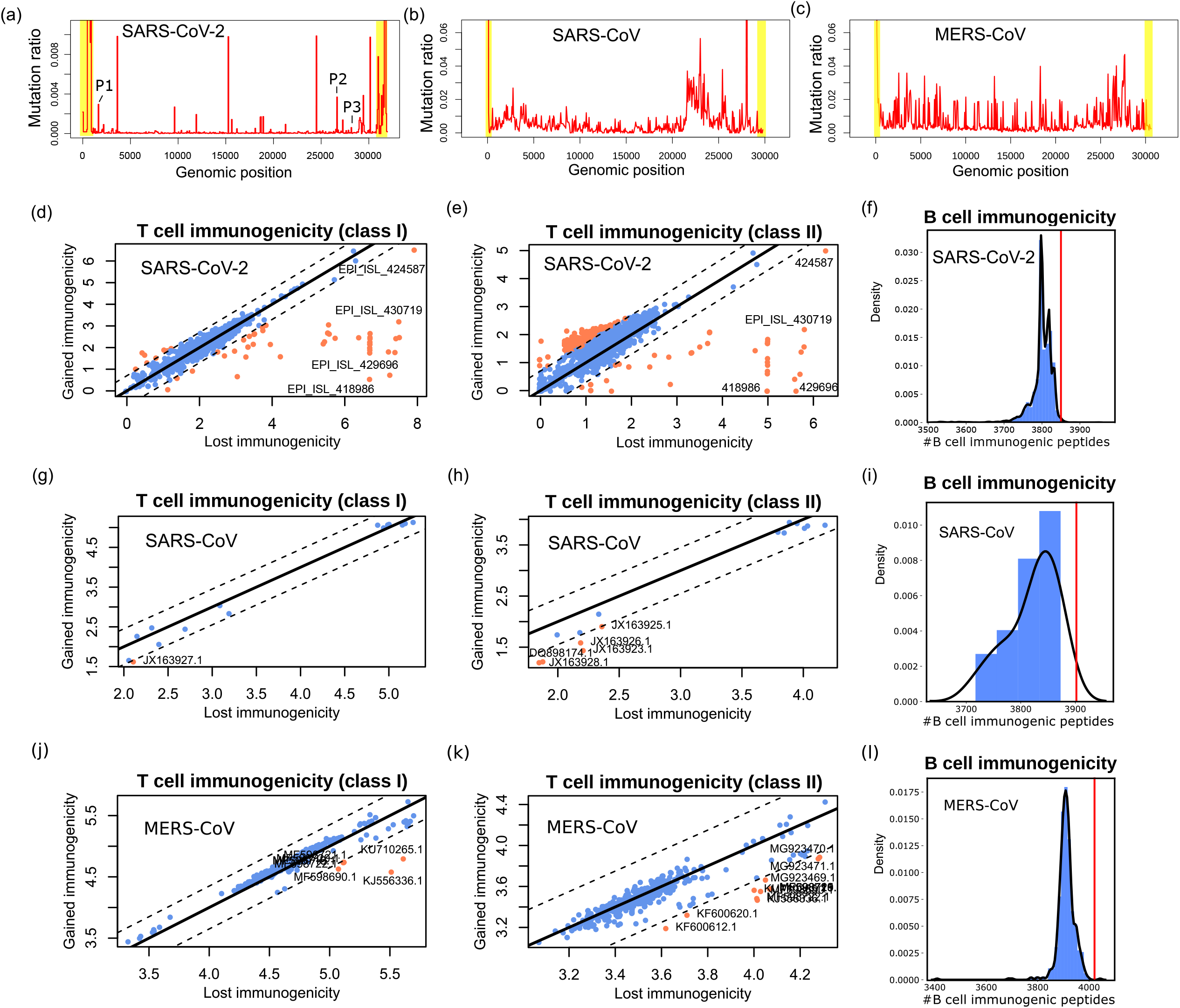
Genetic drifts in SARS-CoV-2 are associated with changes in immunogenicity changes. (a-c) The relative mutational profiles of the three coronaviruses: (a) SARS-CoV-2, (b) SARS-CoV, and (c) MERS-CoV. The Y axis shows the percentage of nucleotides (nts) from all strains of each virus (n=5,568 for SARS-CoV-2, 19 for SARS-CoV-, and 519 for MERS-CoV) in each 60-nt bin that are different from the reference genome. The semi-transparent boxes mark the regions of high mutationals due to artefacts of incomplete sequencing. These regions are shielded from calculations of gain and loss of immunogenicity. The P1-3 labels in **Fig. 2a** refer to the genomic positions with high levels of mutations that have also caused dramatic increases in CD4^+^ T cell immunogenicity (**Fig. 3e**). (d-f) The “lost” immunogenicity and “gained” immunogenicity due to mutations in each SARS-CoV-2 isolate compared with the reference genome. (d) CD8^+^ T epitopes, (e) CD4^+^ T epitopes, and (f) B cell epitopes. The red line indicates the total number of predicted B cell epitope-encoding amino acids in the reference sequence. (g-i) The same immunogenicity change analyses as in (d-f), but for SARS-CoV. (j-l) The same analysis for MERS-CoV.

Next, we examined the potential of the viral proteins to encode linear B-cell epitopes that can elicit antibody responses using BepiPred 2.0 (Jespersen et al., 2017). We focused on linear epitopes, rather than conformational epitopes, because bioinformatics predictions are more feasible for linear epitopes. Contiguous stretches of >10 epitope-encoding residues, which are more representative of the usual length of B cell epitopes, are marked as red bars in **Fig. 1c**, and shown in **Sup. Table 2** in bins of 60 nucleotides. We also examined the receptor-binding motif (RBM) of the SARS-CoV-2 S protein, which attaches to the ACE-2 protein for entry into the human cell (Wan et al., 2020). We used BLAST to align the RBM sequence of the SARS-CoV-2 S protein with that of SARS-CoV, and found there is relatively poor conservation between the two S proteins (**Fig. 1d**). This was supported by Wrapp *et al* (Wrapp et al., 2020), which reports no binding to the SARS-CoV-2 RBM by any of the three SARS-CoV RBM-directed antibodies.

For comparison, we also computed the immune vulnerability maps of SARS-CoV (**Fig. 1e**) and MERS-CoV (**Fig. 1f**), which are the two other coronaviruses known to have caused past outbreaks. We found that the B cell epitope profiles seem to be more consistent among the three viruses. For example, they all share large numbers of predicted B cell epitopes towards the N proteins, which is consistent with the fact that antibody response in SARS-CoV patients is directed most frequently to the N protein (Flego et al., 2005). In contrast, the T cell epitope profiles are more distinct. For example, SARS-CoV Orf1ab protein uniquely possesses many T cell epitopes.

### Genetic drifts of SARS-CoV-2 are associated with immunogenicity changes of the viral proteins

We examined the variants in the viral genomes of SARS-CoV-2, SARS-CoV, and MERS-CoV (**Fig. 2a-c**). SARS-CoV-2 has a much lower level of genetic variation (**Fig. 2a**, n=5,568). In comparison, SARS-CoV (**Fig. 2b**, n=19) and MERS-CoV (**Fig. 2c**, n=519) have both accumulated significant variation among the different strains. MERS-CoV has the highest level of genetic variations as it has spilled over many times into humans over several years. SARS-CoV-2 and SARS-CoV have similar time spans of circulation in humans, and SARS-CoV-2 is still much more slowly mutating than SARS-CoV. In spite of this, in SARS-CoV-2, narrow domains of genomes that are mutated are already discernible, while mutations were observed in other regions as well (**Fig. 2a**). Additionally, more mutations are expected in the viral genome in the future, especially given the possibility that SARS-CoV-2 could continue to propagate in human populations for years to come.

Genetic variations may modify the immunogenicity landscape of the virus. The selection of effective vaccination epitopes should focus on parts of viral proteins with high potential for generating immunogenic epitopes, and with less chance of mutation. We examined whether the genetic drifts of SARS-CoV-2 are associated with the changes in the immunogenicity landscape of this virus (whether some of them are non-random antigenic drifts), and compared against SARS-CoV and MERS-CoV. We examined individual mutations in each of 5,568 SARV-CoV-2 isolates with respect to the reference genome. For each virus isolate, we also calculated the T cell epitopes and B cell epitopes that would be lost and gained due to the mutations in each strain compared to the reference virus genome. For T cell epitopes, we take a sum of the lost/gained epitopes weighted by the Chinese HLA allele frequency, to generate an overall “lost” (compare each strain against the reference) immunogenicity score, and an overall “gained” immunogenicity score. **Fig. 2d** (CD8^+^ T epitopes) and **Fig. 2e** (CD4^+^ T epitopes) show that the gained and lost immunogenicity scores are very comparable for most strains, which is just an effect of random change in the epitope profiles due to mutations. However, for epitopes of CD8^+^ T cells, which are the major cytotoxic T cell population, a small number of strains that showed a large difference in the lost and gained immunogenicity mainly have net loss, rather than net gain, of immunogenicity (**Fig. 2d**). In contrast, the epitopes of CD4^+^ T cells, which have much more complicated biological functions, have an intriguing increase in immunogenicity in a large number of strains, while also having a small number of strains demonstrating net loss in immunogenicity.

Similarly, we calculated the number of amino acid residues that could encode B cell epitopes (**Fig. 2f**). Despite that neutralizing antibody-based vaccines are usually designed against viral surface proteins, there are reports that viral interior and non-structural proteins could also elicit protective humoral immune effects (Carragher et al., 2008; Chen et al., 2018; Liu et al., 2018). Therefore, we considered B cell epitopes of all SARS-CoV-2 proteins and found that there is a dramatic overall decrease in the number of B cell epitopes in the isolates compared with the reference genome (Pval<1e-15). We limited this analysis to amino acid residues in stretches of >10 epitope-encoding residues and had a similar observation (Pval<1e-15, **Sup. Fig. 3a**). These results suggest the existence of an association between humoral immunity and the genetic drifts of SARS-CoV-2 at this stage of human circulation.

**Fig. 3.**
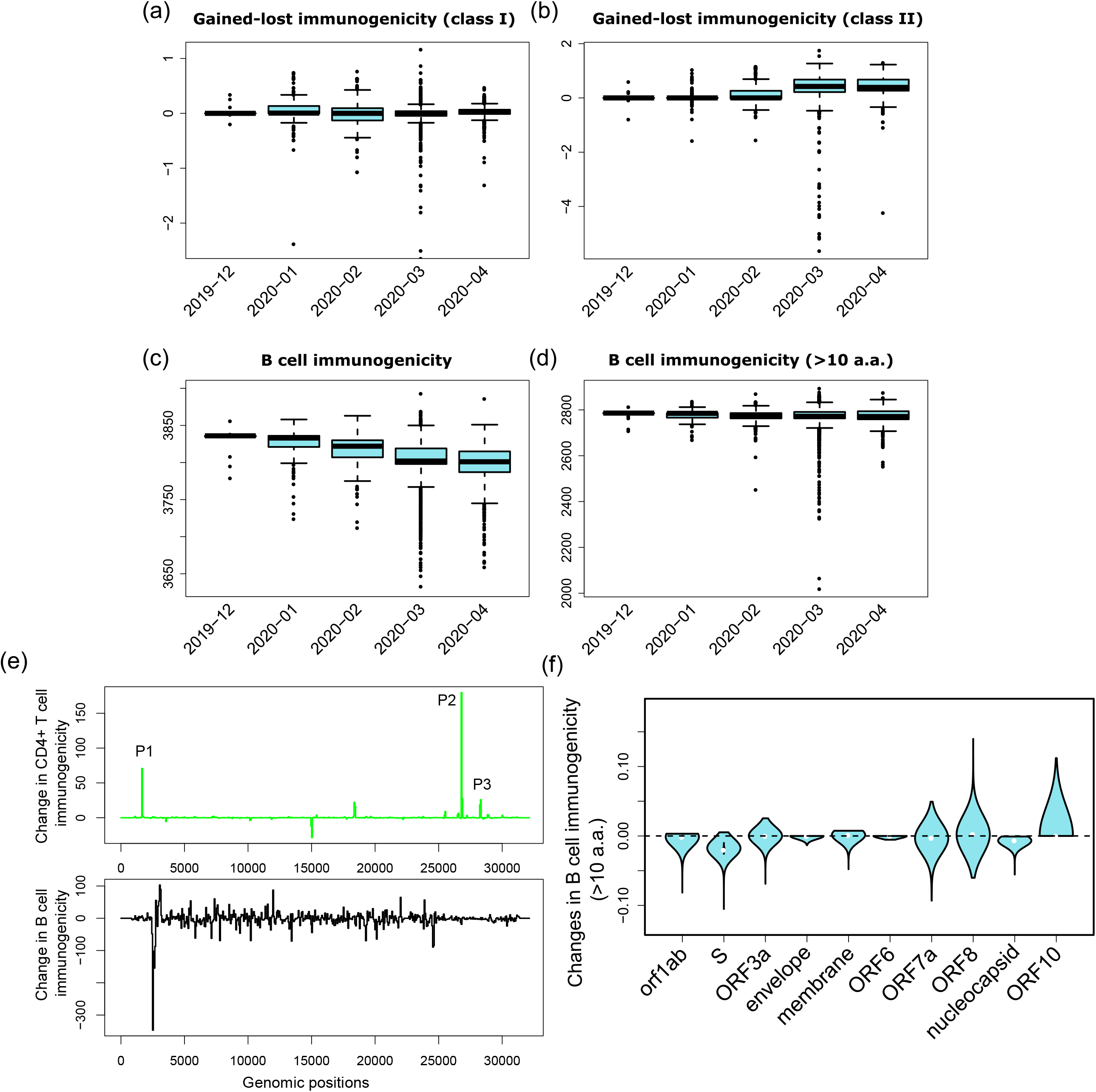
The trend of immunogenicity changes of SARS-CoV-2 over time. (a) Gained minus lost immunogenicity (class I), (b) Gained minus lost immunogenicity (class II), (c) B cell immunogenicity (all epitope-encoding residues), and (d) B cell immunogenicity (epitope-encoding residues in stretches of >10 epitope-encoding residues). The strains were grouped according to their collection time of Dec/2019, Jan/2020, Feb/2020, March/2020, and April/2020. Medians for each group were listed below the plots. (d) The rate of B cell immunogenicity loss in each SARS-CoV-2 viral protein. We calculated, for each viral strain, how many B cell epitopes (>10 a.a.) were lost in each protein. Then we normalized the number of lost epitopes by the length of each protein. (e) The gained/lost predicted CD4^+^ T cell epitopes and B cell epitopes on each genomic position of the SARS-CoV-2 genome across all strains analyzed. The P1-3 labels refer to the genomic positions with dramatic increases in CD4^+^ T cell immunogenicity that also correspond to high levels of mutations (**Fig. 2a**). (f) The rate of B cell epitope on each of the SARS-CoV-2 proteins. The B cell epitope changes were averaged across all viral strains, for each protein, and then normalized by the length of that protein.

In comparison, we performed a similar association analysis between genetic variation and immunogenicity for SARS-CoV (**Fig. 2g-i** and **Sup. Fig. 3b**) and MERS-CoV (**Fig. 2j-l** and **Sup. Fig. 3c**). In contrast to SARS-CoV-2, SARS-CoV and MERS-CoV strains with large immunogenicity changes almost all have a net loss, rather than net gain, of immunogenicity, for both CD8^+^/CD4^+^ T cell epitopes and B cell epitopes.

Furthermore, we grouped the SARS-CoV-2 strains by their collection times (Dec/2019, Jan/2020, Feb/2020, March/2020, and April/2020), and compared the changes in T cell and B cell immunogenicity. This longitudinal analysis shows that the number of CD8^+^ T cell epitopes showed no clear trend over time, and the latter appearing strains have gained immunogenicity for CD4^+^ T cells (**Fig. 3ab**, Jonckheere-Terpstrata test Pval=4.657e-11). The same analyses regarding T cell epitopes weighted by the Caucasian, Black, Latino, and Middle Eastern allele frequencies resulted in very similar conclusions (**Sup. Fig. 4**). Consistent with our observations above, **Fig. 3c** shows the latter appearing strains are less immunogenic than earlier strains with statistical significance achieved for B cell epitopes (Jonckheere-Terpstrata test Pval<2.2e-16). We observed the same trend when focusing on B cell epitopes that are in stretches longer than 10 epitope-encoding amino acids (**Fig. 3d**).

**Fig. 4.**
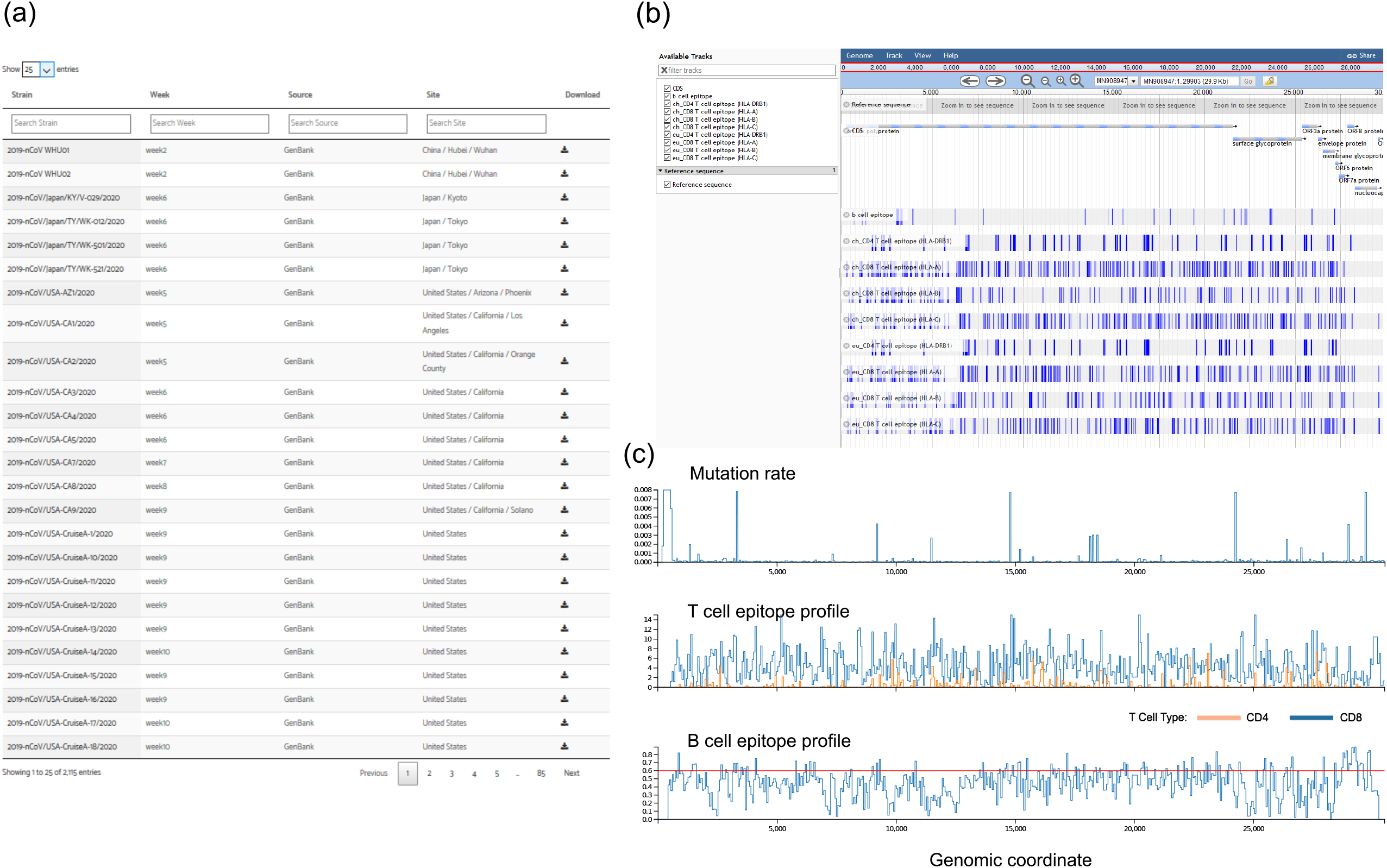
An actively updated database of the immune vulnerability landscape of SARS-CoV-2. (a) A searchable table of all the 5,568 SARS-CoV-2 strains. The strain names and relevant annotation information are provided in the table. (b) Upon clicking the download button, the user will be able to download a small data packet for the corresponding viral strain, which can be visualized in JBrowse desktop (https://jbrowse.org/blog). A screenshot of an example JBrowse session is shown. (c) Additional visualization functionality to examine immunogenicity profiles of the selected genomic region of the viruses (the Chinese population). Each line plot is divided into two panels stacked vertically together. At the bottom panel, the user can drag and set a region to zoom in, and the top panel zooms in and shows the details of that selected region.

Next, we investigated the genomic locations of the antigen variants. We examined all genomic positions of all SARS-CoV-2 strains and counted the number of gained/lost B cell and CD4^+^ T cell epitopes. Interestingly, for CD4^+^ T cell epitopes, the genomic positions with large net gains in immunogenic epitopes are concentrated in several sharp peaks (P1-P3 of **Fig. 3e**), and these sharp peaks correspond to genomic positions with high levels of mutations in SARS-CoV-2 (P1-P3 of **Fig. 2a**). Upon closer examination, there is one driving mutation within each peak that accounts for most of the mutational events, but there are also many low allele frequency mutations as well. We predicted T_reg_ epitopes using the EpiVax JanusMatrix algorithm (Moise et al., 2013), which successfully identified HCV and flu T_reg_ epitopes (Liu et al., 2015; Losikoff et al., 2015) and has been applied to develop flu vaccines (Wada et al., 2017). Interestingly, we found that the driving mutation in P3 has enhanced T_reg_ epitope potential (**Sup. File 1**), which could induce tolerogenicity and immune escape, and might explain why this mutation has been selected for overtime.

In comparison, the distribution of the changed B cell epitopes shows a different pattern. **Fig. 3e** shows that the B cell epitope changes are much more evenly spread out across the whole genome. There is a sharp peak of loss of B cell immunity around 2,500bp on Orf1ab, but immediately 3’ of this position, there is another peak of gain of B cell immunity. Curiously, these two peaks do not correspond to the hotspots of mutations (**Fig. 2a**). Nevertheless, this result suggests this region of Orf1ab might be important for the interaction of SARS-CoV-2 with the host humoral response system, which warrants further investigation. Overall, it seems that the changes in B cell immunogenicity is a general pattern that happens in the whole viral genome. However, we still suspect that there is a difference in the tendency of different viral proteins to lose B cell immunogenicity. We examined each of the SARS-CoV-2 proteins and calculated the rate of B cell immunogenicity loss normalized by the length of each protein across all viral strains (**Fig. 3f**). As is expected, the S protein shows the highest rate of B cell immunogenicity loss, followed by the nucleocapsid protein. Although the antibodies against the nucleocapsid proteins are likely non-neutralizing, Carragher *et al* showed that they could still play important roles in facilitating resistance to viral infection (Carragher et al., 2008), which might explain our observation.

### A actively updated database of the immune vulnerability landscape of SARS-CoV-2

To facilitate immunological studies of SARS-CoV-2, we created the SARS-CoV-2 Immune Viewer (**Fig. 4**) to openly share the viral immunogenicity data of SARS-CoV-2, and also SARS-CoV and MERS-CoV. All SARS-CoV-2 strains that we have downloaded and processed are presented in a searchable table, with annotation information associated with each strain (**Fig. 4a**). Upon clicking each row of the table, users will be able to download a small data packet for that strain of SARS-CoV-2, which can be directly visualized on the desktop JBrowse software (Buels et al., 2016) (https://jbrowse.org/blog/). The JBrowse visualization demonstrates the genomic sequence, protein annotations, and T cell and B cell epitopes of each viral isolate (**Fig. 4b**). Within the Immune Viewer, the densities of the mutations and the T cell and B cell epitopes are also shown in interactive plots (**Fig. 4c**). This resource will be updated roughly monthly as new SARS-CoV-2 strains become available (with 1-2 months needed to process the new batch of viral sequences collected in each month).

## DISCUSSION

In summary, this study provided the immune vulnerability landscape of SARS-CoV-2 and investigated the association between the genetic drifts of SARS-CoV-2 and immunogenicity changes of the viral proteins by analyzing more than 5,000 virus genomes. We also compared the findings from SARS-CoV-2 with those of SARS-CoV and MERS-CoV. This work is complementary to the work of Grifoni *et al*, where a sequence homology approach together with bioinformatics analysis was taken to map SARS-CoV-2 proteins to experimentally validated immunogenic epitopes (Grifoni et al., 2020). However, that work only used the reference genome of the SARS-CoV-2 virus and did not analyze the genetic drifts and its association with immunogenicity. Our work, for the first time, clearly demonstrated the evolution of immunogenicity in SARS-CoV-2, and also compared it with the evolution of SARS-CoV and MERS-CoV. To disseminate our research, a publicly accessible database, the SARV-CoV-2 Immune Viewer, was developed for easy exploration and downloading of results. The database is under continuous maintenance and updated roughly monthly to incorporate new strains of SARS-CoV-2 made available. We believe this is the first public immune-profiling resource of SARS-CoV-2 (including all its strains) for the research community.

This study shows that the mutations in SARS-CoV-2 could be more than merely random genetic drifts, by showing that some strains of SARS-CoV-2 have immunogenicity loss of B cell-based host immune surveillance and gain of CD4^+^ T cell immunogenicity due to mutations. A minimal loss in CD8^+^ cell-based immunogenicity was also observed. Partially similar to SARS-CoV-2, SARS-CoV and MERS-CoV have obvious loss of both CD8^+^ T cell-based and B cell-based immunogenicity. On the other hand, it is very intriguing that CD4^+^ T cell-based immunogenicity increased in SARS-CoV-2, whereas decreases were observed in SARS-CoV and MERS-CoV. Our analyses provided one possible explanation that, for at least the driving mutation of P3, the increased CD4^+^ immunogenicity could actually be an increase in the potential of the epitope to induce T_reg_ response, which might confer a survival advantage to the virus.

The loss of immunogenicity will lead to a loss of resistance to viral infection. However, it is uncertain whether the loss of immunogenicity will lead to a worsening of symptoms of the patients when infected, which is dependent on complicated biological and immunological processes in the human body. It is possible that a virus will become more infectious, but also less pathogenic at the same time, which may bring a stronger survival advantage to the virus. This effect is likely hard to investigate through simplistic testing of diversifying selection, such as the observed versus expected dN/dS substitution ratios, but solving this paradox could be particularly important for tackling the outbreak of SARS-CoV-2. This effect may also be related to a large number of reported asymptomatic cases of SARS-CoV-2 (Ling et al., 2020; Pan et al., 2020), and the observation that SARS-CoV and MERS-CoV are considerably more aggressive than SARS-CoV-2 in terms of mortality rates.

Admittedly, bioinformatics prediction of B cell epitopes is still challenging (Galanis et al., 2019). However, the Bepipred 2.0 software that we used is among the best such software. More importantly, our conclusions regarding immunogenicity loss or gain are drawn based on the overall trend of changes in the total number of predicted B (and similarly T) cell epitopes among different strains. This analysis aggregates individual predictions with random and independent errors and is thus subject to much less inaccuracies.

Overall, this study provides a window into the immunological features of SARS-CoV-2, and have yielded curious insights into the evolution of this virus. This resource may aid therapeutic and vaccination development against this virus to stop this pandemic earlier and to prevent future outbreaks.

## MATERIALS AND METHODS

### Acquisition of the viral genome sequences

The SARS-CoV-2 complete genome sequences and meta data were downloaded from the NGDC (https://bigd.big.ac.cn/ncov) and GISAID (https://www.gisaid.org/) databases. Sequences that are duplicated (8.7% of all collected strains), as defined by sharing the same genomic sequences, or that have low coverage (12.9%) were not included in the analyzed strains. The rest analyzed strains include all those submitted before April 6^th^ and 3,454 strains collected between April 6^th^ and May 13^th^, which were randomly sampled from all available strains in that time window. This down-sampling was done as the sequence alignment became impractically slow if all strains were included. The reference genome was acquired from NCBI, which is one of the first few isolates of SARS-CoV-2 collected and sequenced in late December of 2019: https://www.ncbi.nlm.nih.gov/nuccore/MN908947.

The removal of the duplicates alleviates the concern in potential redundancy of the genomic data imposed by a biased sampling. Admittedly, left-over bias in sampling could still exist in the sampled strains, even after duplicate removal. However, it is difficult to further adjust for this bias phylogenetically, which could be performed through certain sub-sampling approaches with respect to the size of the infected population in each country. However, the reported numbers of infection cases in different countries are subject to inaccuracies due to methods of testing and other reasons, making this approach likely to result in additional bias.

The complete genome SARS-CoV and MERS-CoV sequences are also downloaded from NCBI: https://www.ncbi.nlm.nih.gov/nuccore/?term=txid694009%5BOrganism%3Anoexp%5D+and+complete+genome and https://www.ncbi.nlm.nih.gov/nuccore/?term=txid1335626%5BOrganism%3Anoexp%5D+and+complete+genome. The reference genomes (SARS-CoV: NC_004718.3 and MERS-CoV: NC_019843.3) were determined by NCBI, which were one of the earliest few sampled isolates for each coronavirus, respectively (sometimes only the months of sample collection were shown). Each isolate’s genomic sequence was aligned to the reference genome sequence using EMBOSS needle with the gap opening penalty of 20 and the gap extension penalty of 0. The sequence differences were identified as genomic variants and they were annotated using Annomen. In **Fig. 2a-c**, the semi-transparent boxes mark the regions of high mutations due to artefacts of incomplete sequencing. These regions are shielded from calculations of genomic variants. The protein sequences of the isolates were determined based on the sequence differences and the reference gene annotations. The isolate protein sequences were then aligned to the reference protein sequences using EMBOSS needle and the strains whose protein alignments do not cover >90% of the reference protein sequence were ignored.

### DNA and protein sequence alignment

The command-line version of MUSCLE (v3.8.31) (Edgar, 2004a, 2004b) was used to perform multiple genome sequence alignment with diagonal optimization (-diags). The default number of iteration and the default maximum number of new trees were applied during the alignment. The protein sequence alignment between the S proteins, YP_009724390.1 (SARS-CoV-2) and NP_828851.1 (SARS-CoV), was performed using EMBOSS needle(Rice et al., 2000) with the BLOSUM62 scoring matrix.

### Prediction of T cell and B cell epitopes

NetMHCpan (v4.0) (Jurtz et al., 2017) and NetMHCIIpan (v3.2) (Jensen et al., 2018) with default threshold options were used to predict T cell peptides from the viral proteins that bind to human MHC class I and II proteins for all the available HLA alleles. The HLA allele population frequency for the Chinese population was acquired from Kwok *et al* (*Kwok et al*., *2016*) and population frequencies for the U.S. European Caucasian, African American, Hispanic (Central and South America), and Middle East/North African populations were obtained from the Allele Frequency Net Database, originally from Gragert *et al* (González-Galarza et al., 2015). The B cell epitope predictions were made by the BepiPred 2.0 software (Jespersen et al., 2017), with default parameters. Amino acids with B cell epitope prediction scores >0.6 are regarded as having high likelihoods of generating linear antibodies.

### Website development

The Immune Viewer is a dynamic website. It is developed using HTML (HyperText Markup Language), JavaScript and CSS (Cascading Style Sheets). Specifically, we used the D3.js library to allow users to interactively explore the mutations or immunogenic scores across the viral genomic regions. We also used the D3.phylogram.js to visualize the phylogenetic tree and the Select2 library to facilitate users’ query for different SARS-CoV-2 strains across multiple geographic regions. The data packet downloaded for each viral strain can be directly visualized in the desktop JBrowse software, which can be downloaded from https://jbrowse.org/blog.

### Statistical analyses

All computations and statistical analyses were carried out in the R and Python computing environment. For all boxplots appearing in this study, box boundaries represent interquartile ranges, whiskers extend to the most extreme data point which is no more than 1.5 times the interquartile range, and the line in the middle of the box represents the median. For the line plots, the viral genomes were binned by every 60 nucleotides, and the number of T cell and B cell epitopes within each window is calculated. For T cell epitopes, a sum of the number of epitopes weighted by the corresponding ethnic population’s HLA allele (A, B, C, and DRB1) frequency is calculated to form the T cell immunogenicity strength for that population. The genetic variation ratio at each nucleotide is calculated by examining all viral strains and counting the proportion of strains with a different nucleotide or with an insertion/deletion, with respect to the reference genome. The genetic variation ratios are also binned by the same length of windows and averaged.

**Sup. Fig. 1.**
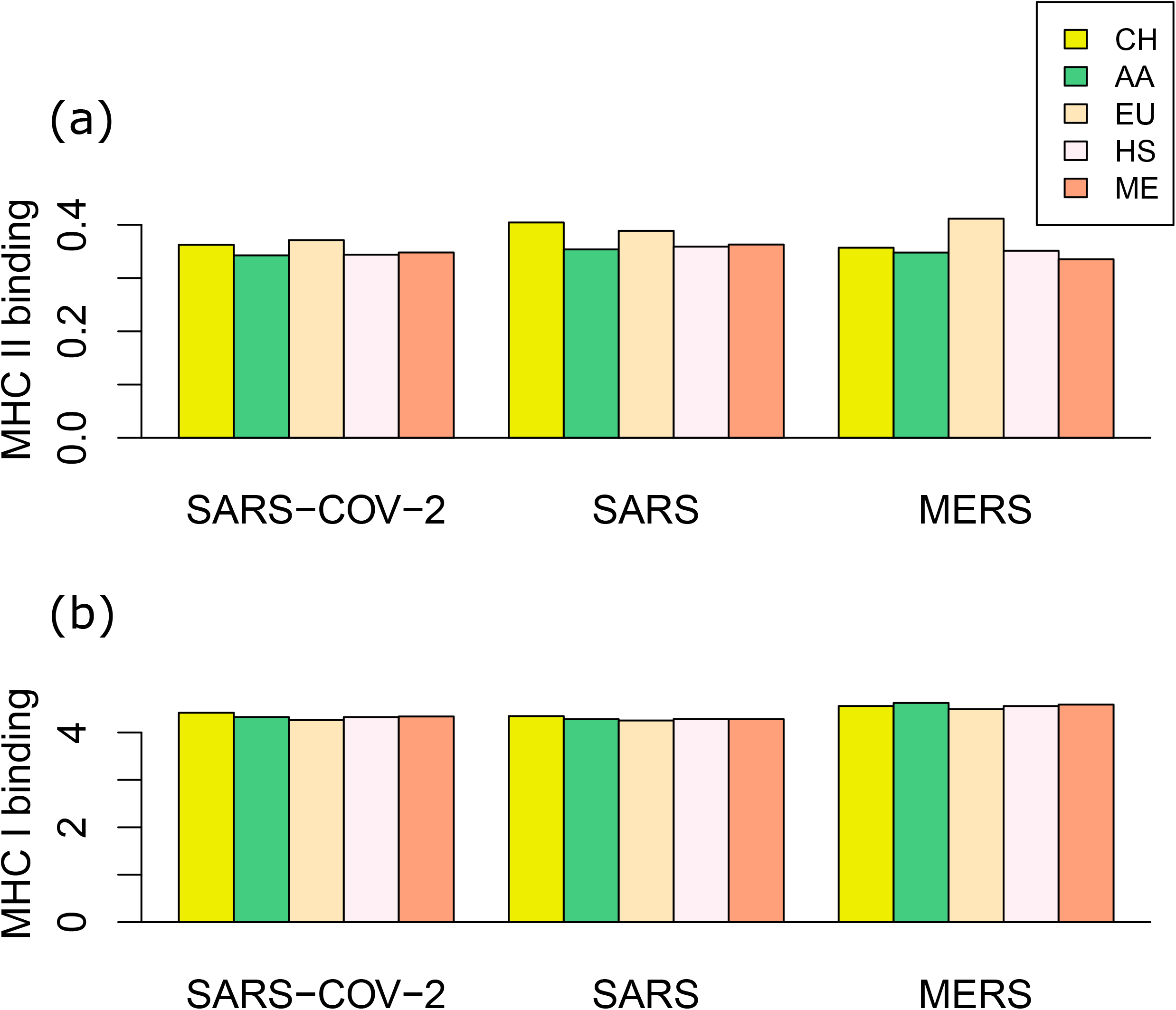
The overall T cell immunogenicity strengths of SARS-CoV-2, SARS-CoV, and MERS-CoV in several racial populations. (a) predicted CD4^+^ T cell immunogenicity; (b) predicted CD8^+^ T cell immunogenicity. The T cell immunogenicity density scores in all bins of each genome, as defined in **Fig. 1a**, are averaged. (CH: Chinese, AA: African American, EU: European Caucasian (U.S.), HS: Hispanic (Central and South American, U.S.), ME: Middle East/North African (U.S.)).

**Sup. Fig. 2.**
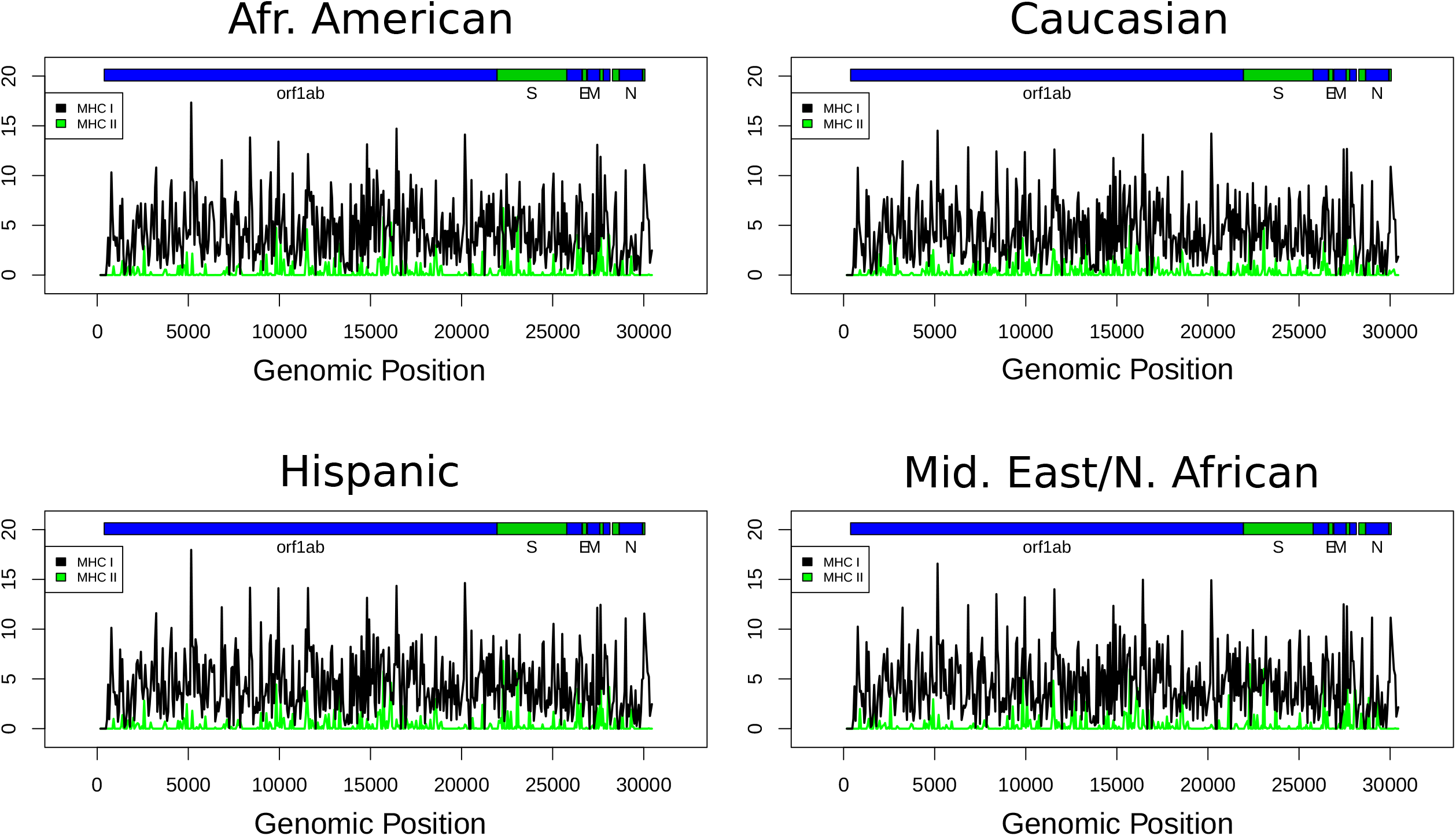
The T cell epitope profiles of SARS-CoV-2, in the African American, Caucasian, Hispanic, and Middle Eastern/North African populations.

**Sup. Fig. 3.**
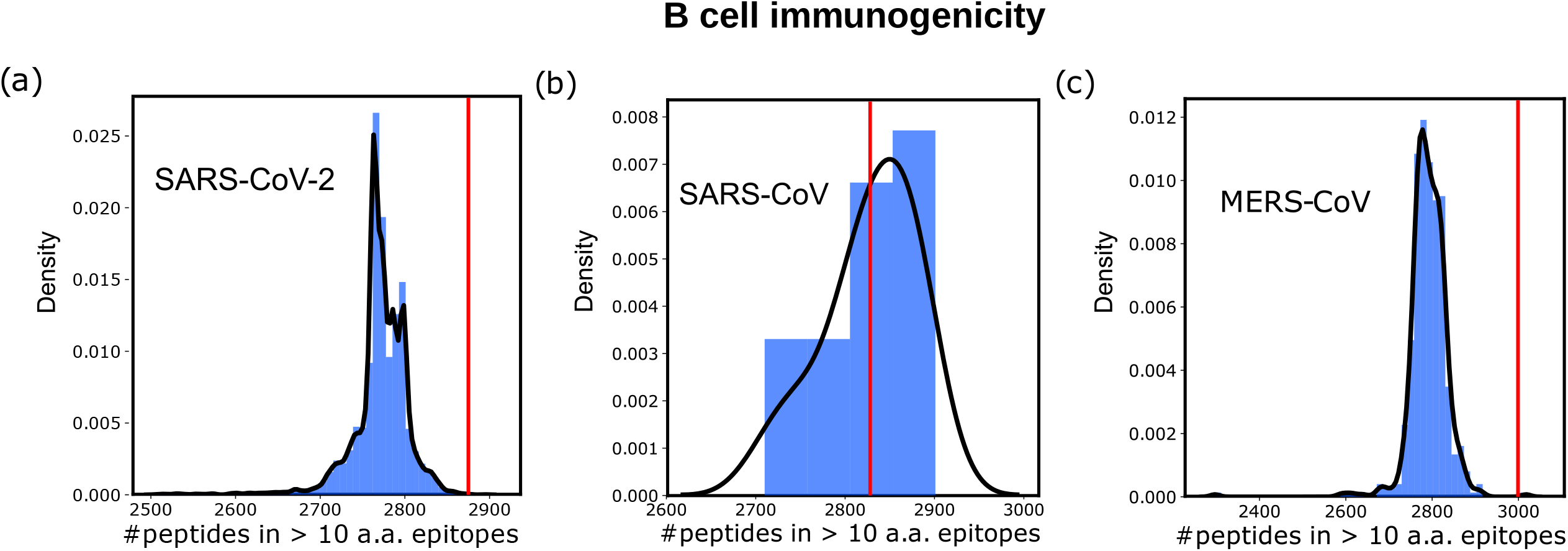
B cell epitope-encoding amino acids in contiguous stretches of >10 epitope-encoding amino acids. (a) SARS-CoV-2, (b) SARS-CoV, and (c) MERS-CoV. Red lines denote the total numbers of epitope-encoding amino acids of the reference sequences.

**Sup. Fig. 4.**
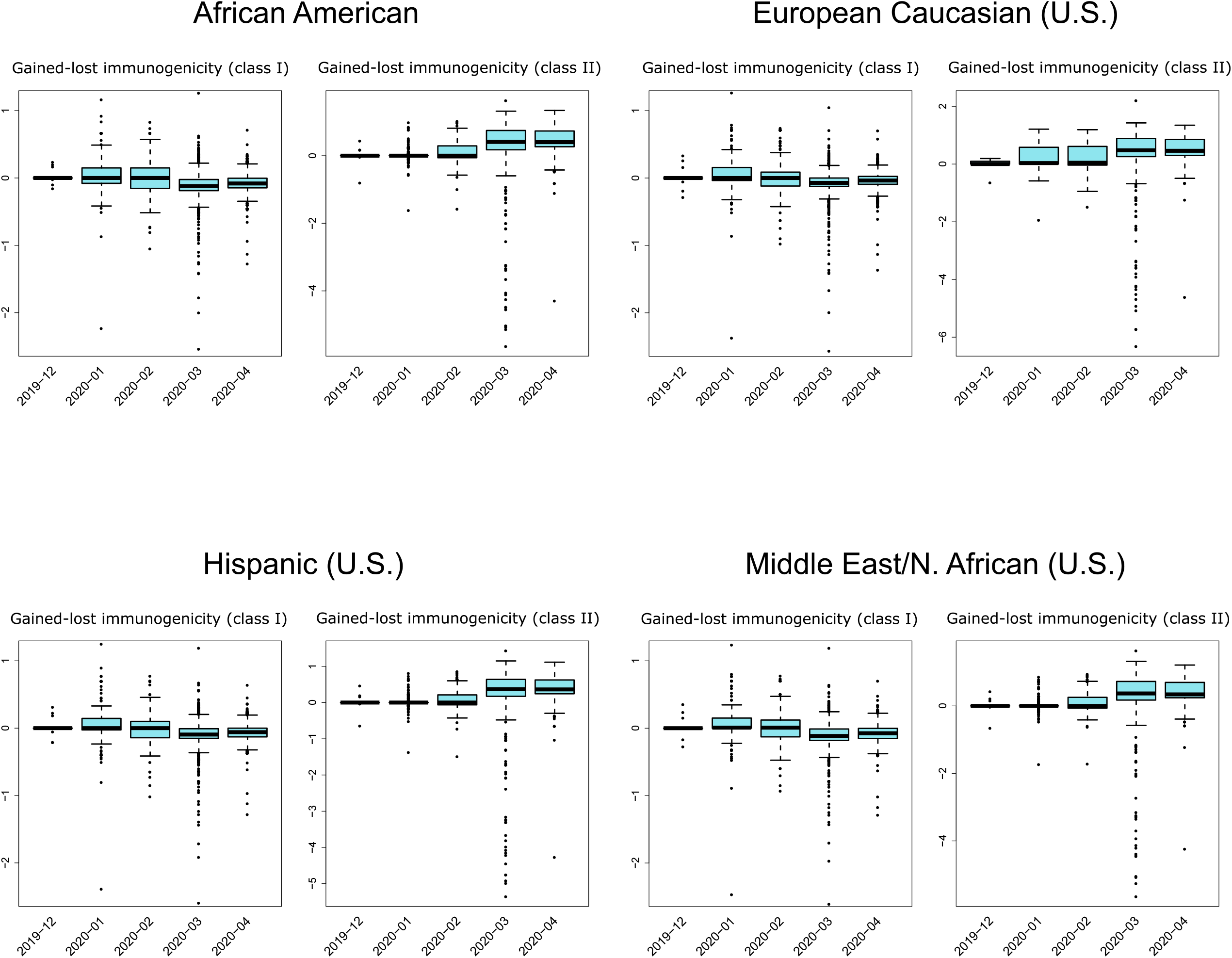
The trend of T cell immunogenicity changes of SARS-CoV-2 over time, with T cell immunogenicity weighted by the HLA allele frequencies of the Caucasian, Black, Latino, and Middle Eastern populations.

**Sup. Table 1** Genomics regions of SARS-CoV-2 that are T cell epitope-enriched. The top 50 regions with the highest predicted binding were shown, for CD4^+^ and CD8^+^ T cells, respectively. IEDB records of experimental validation are also shown in the table.

**Sup. Table 2** Genomics regions of SARS-CoV-2 that are B cell epitope-enriched. The top 91 regions with predicted B cell epitope scores > 0.6 are shown.

**Sup. File 1** EpiVax JanusMatrix prediction results for the mutations in P1-3.

## DATA AVAILABILITY

The SARS-CoV-2 Immune Viewer is publicly accessible, with the genome sequence, protein annotation, T/B cell epitope prediction results freely downloadable for each strain: https://qbrc.swmed.edu/projects/2019ncov_immuneviewer/.

## FUNDING

This study was supported by the National Institute of Health [1R01GM115473-01A1, 5P30CA142543-07] and Cancer Prevention Research Institute of Texas [CPRIT RP190208, and RP180805]

### ACKNOWLEDGEMENTS

We acknowledge the patients who contributed the viral islets, the medical staff and researchers for performing islet purification and sequencing, and the NGDC (https://bigd.big.ac.cn/ncov) and GISAID (https://www.gisaid.org/) databases for timely sharing of the viral genome sequences. We acknowledge Drs. Annie De Groot, Leonard Moise, and Andres Gutierrez for performing the JanusMatrix analysis.

## AUTHOR CONTRIBUTIONS

J.Z. and X.Z. performed statistical analyses of the immune vulnerability landscape of the viruses. J.K., T.W., X.X, and X.Z. retrieved and curated the virus sequence genomes. J.K. and J.Z. performed T cell and B cell epitope predictions. X.X. and J.K carried out DNA and protein sequence alignment. Y.W., D.L., R.C., and X.Z. created the Immune Viewer. H.Z. and X.Z. calculated the phylogenetic trees for the website. J.S., G.X. and L.X. provided significant input on the scientific direction of the project. T.W. and Y.X. supervised the study. J.Z., J.K., X.X., Y.W., X.Z., T.W., and Y.X wrote the manuscript.

