## Supplementary material for "The immune vulnerability landscape of the 2019 Novel Coronavirus, SARS-CoV-2": Sup. File 1

**The modified peptides (Mod) shown below have the most significant changes compared to the original T cell epitope clusters from P1, P2 and P3 of the reference SARS-CoV-2 genome. The driving mutations in P1-3 are highlighted in yellow, which account for most of the mutational events in each peak.**

**EMX woFlanks: predicted immunogenicity by the EpiVax EpiMatrix algorithm (not the netMHCIIpan algorithm). “Green” shows gain and “red” shows loss. One of the driving mutations shows a significant gain in immunogenicity.**

**JMX (JanusMatrix): predicted T_reg_ epitope potential. “Red” shows gain in T_reg_ tolerogenicity potential, and “green” shows loss. The driving mutation in P3 shows a significant gain in the predicted potential of becoming a T_reg_ epitope.**

```
>NSP2 (P1)
AYTRYLIGGVDNNFCGPDGYPLECIKDLLARAGKASCTLSEQLDFIDTKRGVYCCR
EHEHEIAWYTERSEKSYELQTPFEIKLAKKFDTFNGECPNFVFPLNSIIKTIQPRV…

Reference genome T cell epitope clusters                                           EMX woFlanks,JMX
72-88              QTPFEIKLAKKFDTFNG                                                     18.15,1.10
83-97                         FDTFNGECPNFVFPL                                            13.17,0.67
89-108                              ECPNFVFPLNSIIKTIQPRV                                 18.19,1.33

Modifications that may impact T cell epitope clusters
72_Mod1            QTPFEIKLAKKFDIFNG                                                     21.44,1.09
83_Mod1                       FDTFNEECPNFVFPL                                            12.12,0.00
89_Mod1                             ECPNFVFSLNSIIKTIQPRV                                 41.30,1.10


>ORF3a QHD43417 (P2)
…VRATATIPIQASLPFGWLIVGVALLAVFQSASKIITLKKRWQLALSKGVHFVCNLLLLFVTVYSHLLLVAAGLEAPFL…

Reference genome T cell epitope clusters                                           EMX woFlanks,JMX
42-58         PFGWLIVGVALLAVFQS                                                          24.28,4.23
53-67                    LAVFQSASKIITLKK                                                 26.09,0.00
67-82                                  KRWQLALSKGVHFVCN                                  23.09,2.42
74-106                                        SKGVHFVCNLLLLFVTVYSHLLLVAAGLEAPFL          95.68,3.26

Modifications that may impact T cell epitope clusters
42_Mod1       PFGWLIVGVAFLAVFQS                                                          11.93,0.75
42_Mod2a      PFGWLIVGVALLVVFQS                                                          24.12,2.79
42_Mod2b      PFGWLIVGVALLSVFQS                                                          30.18,3.11
42_Mod3       PFGWLIVGVALLAGFQS                                                          22.31,3.33
42_Mod4       PFGWLIVGVALLAVCQS                                                          24.28,4.23
42_Mod5       PFGWLIVGVALLAVFHS                                                          24.28,4.00

53_Mod1a                 LVVFQSASKIITLKK                                                 26.09,0.00
53_Mod1b                 LSVFQSASKIITLKK                                                 26.09,0.26
53_Mod2                  LAGFQSASKIITLKK                                                 26.09,0.13
53_Mod3                  LAVCQSASKIITLKK                                                  4.40,0.00
53_Mod4a                 LAVFHSASKIITLKK                                                 28.50,0.19
53_Mod4b_2Mut            LAVFHSASKIITFKK                                                 28.50,0.19
53_Mod5a                 LAVFQSDSKIITLKK                                                 10.15,0.00
53_Mod5b                 LAVFQSVSKIITLKK                                                 25.44,0.00
53_Mod6                  LAVFQSASNIITLKK                                                 30.07,0.00
53_Mod7                  LAVFQSASKIITFKK                                                 26.09,0.00
53_Mod8                  LAVFQSASKIITLTK                                                 26.09,0.00
53_Mod9                  LAVFQSASKIITLKN                                                 26.09,0.00

67_Mod1                                NRWQLALSKGVHFVCN                                  23.09,2.42
67_Mod2                                KRLQLALSKGVHFVCN                                  21.41,4.53
67_Mod3                                KRWQLSLSKGVHFVCN                                  21.29,2.58
67_Mod4a                               KRWQLALFKGVHFVCN                                   4.29,2.25
67_Mod4b                               KRWQLALPKGVHFVCN                                  14.56,3.29
67_Mod5                                KRWQLALSEGVHFVCN                                  14.46,0.38
67_Mod6                                KRWQLALSKSVHFVCN                                  29.03,0.93
67_Mod7                                KRWQLALSKGIHFVCN                                  23.13,2.91
67_Mod8a                               KRWQLALSKGVYFVCN                                  19.42,3.00
67_Mod8b                               KRWQLALSKGVRFVCN                                  21.33,2.70
67_Mod9                                KRWQLALSKGHVFICN                                  16.71,2.88

74_Mod1a                                      FKGVHFVCNLLLLFVTVYSHLLLVAAGLEAPFL          95.68,3.26
74_Mod1b                                      PKGVHFVCNLLLLFVTVYSHLLLVAAGLEAPFL          95.68,3.26
74_Mod2                                       SEGVHFVCNLLLLFVTVYSHLLLVAAGLEAPFL          95.68,3.26
74_Mod3                                       SKSVHFVCNLLLLFVTVYSHLLLVAAGLEAPFL          95.68,3.26
74_Mod4                                       SKGIHFVCNLLLLFVTVYSHLLLVAAGLEAPFL          97.10,3.26
74_Mod5a                                      SKGVYFVCNLLLLFVTVYSHLLLVAAGLEAPFL          93.50,3.26
74_Mod5b                                      SKGVRFVCNLLLLFVTVYSHLLLVAAGLEAPFL          95.96,3.37
74_Mod6                                       SKGVHFICNLLLLFVTVYSHLLLVAAGLEAPFL          95.47,3.13
74_Mod7                                       SKGVHFVCNLLLLFATVYSHLLLVAAGLEAPFL          83.77,3.45


>MEMBRANE QHD434193 (P3)
MADSNGTITVEELKKLLEQWNLVIGFLFLTWICLLQFAYANRNRFLYIIKLIFLWLLWPVTLACFVLAAV
YRINWITGGIAIAMACLVGLMWLSYFIASFRLFARTRSMWSFNPETNILLNVPLHGTILTRPLLESELVI
GAVILRGHLRIAGHHLGRCDIKDLPKEITVATSRTLSYYKLGASQRVAGDSGFAAYSRYRIGNYKLNTDH
SSSSDNIALLVQ…

Reference genome T cell epitope clusters                                           EMX woFlanks,JMX
165-179|Rank 20         PKEITVATSRTLSYY                                                   9.81,0.14
175-190|Rank 10                   TLSYYKLGASQRVAGD                                       17.64,1.83

Modifications that may impact T cell epitope clusters                             EMX woFlanks,JMX
165_Mod1                PKQITVATSRTLSYY                                                   9.81,0.14
165_Mod2                PKEITVATSRMLSYY                                                   9.83,0.00
175_Mod1                          MLSYYKLGASQRVAGD                                       17.64,2.29
```
